# Single-cell western blotting of cytoplasmic cytokeratin 8 proteoforms

**DOI:** 10.1101/2025.01.21.634008

**Authors:** Anna Fomitcheva Khartchenko, Trinh Lam, Amy E. Herr

## Abstract

Differential detergent fractionation (DDF) enables compartment-specific lysis, offering a strategy to analyze cytoplasmic proteins while preserving the nucleus for downstream assays. However, while this method facilitates sub-cellular resolution, current single-cell approaches using DDF remain limited in their ability to identify proteoforms without compromising nuclear integrity. This limitation is especially pronounced in proteins where their proteoforms present diverse biological functions such as cytokeratin 8 (CK8), a structural protein implicated in several disease states. Here, we present a single-cell western blot (scWB) integrated with DDF to selectively solubilize and separate CK8 proteoforms while preserving nuclear integrity. To evaluate assay applicability and nuclear stability, we profiled CK8 across breast cancer cell lines (MCF7, SKBR3, and MDA-MB-231), confirming proteoform detection only in MCF7 and preservation of nuclear content across all lines. We report on assay development, including screening a panel of lysis buffers based on nonionic detergents, and electrophoresis conditions to achieve a separation resolution between two proteoforms of up to 0.94, while preserving an intact nucleus. The cytoplasm-specific lysis (DDF buffer) yielded detectable proteoforms in 14.3% of solubilized cells, comparable to 10.3% with whole-cell lysis (RIPA buffer). Our approach allows for tailored solubilization, achieving reliable proteoform detection and nuclear retention across different cell types. Proteoform profiling at the single-cell level forms a basis for the exploration of the role of specific CK8 molecular forms in cellular processes.

## Introduction

Cytokeratin 8 (CK8) is an intermediate filament located in the cytoplasm and nucleus, that serves as a structural element and mediator of signal transduction. As a type II keratin, CK8 typically dimerizes with CK18^1^, and is involved in various disease states. CK8 expression is modulated during viral infections, such as hepatitis C, influenza, and zika, with evidence suggesting a reorganization of the cytoskeleton to facilitate viral replication^2–4^. As a product of *KRT8* gene mutations, CK8 may form amyloid-like aggregates linked to alcoholic liver disease^5^. Moreover, CK8 and its proteoforms have been implicated in cancer. Panels of cytokeratins have been used for the identification and diagnosis of certain cancer types such as bladder^6^, lung^7^, breast^8^, and head and neck^9^. Notably, a CK8 proteoform in lung adenocarcinoma has been correlated with decreased survival^10^. In colorectal cancer, transcript-level CK8 revealed a clear differentiation of expression between tumor and normal tissue^11^, highlighting the relevance of CK8 as a biomarker.

Currently, UniProt identifies two confirmed alternative-splicing derived isoforms of CK8 (#P05787), as well as six additional shorter forms, some of which are only computationally predicted^12^. At the transcript level, CK8 presents two isoforms. Recently, Li *et al* detected a new splicing variant, which corresponds to a predicted molecular mass of 46.2 kDa^11^, although its translation to protein remains uncertain, as no CK8 fragment with this molecular mass has been reported to our knowledge. The null observation suggests that this isoform is not translated into protein, post-translationally cleaved into shorter forms, or expressed below the detection limit of assays used to interrogate the system. Previous studies provide evidence of CK8 cleaving events, yielding variable-length fragments of this CK8 46.2 kDa form^13^. Getting from transcript to expressed protein presents numerous opportunities for physico-chemical diversity, thus highlighting the need for tools and strategies to analyze the presence of proteoforms as well as additional omics expressions.

To prepare a cell suspension for analyses of targeted cellular compartments, compartment-specific assays have been appended to differential detergent fractionation (DDF) sample preparation approaches. DDF is a chemically targeted approach to selectively lyse cellular organelles, including the cytoplasm, nucleus, and various organelles. DDF mitigates non-specific background interference, commonly arising from cross-contamination between cellular compartments. DDF also offers a cell compartment-specific fractionation of nuclear components for nucleic acid analysis versus cytoplasmic compartments for protein analyses. Bulk DDF methods use differential and density gradient centrifugation and magnetic beads to capture the organelles of interest^14,15^. These bulk approaches, however, mask single-cell heterogeneity. DDF has been adapted to single-cell sample preparation; for instance, single cells isolated in microwells were subjected to selective lysis and electrophoresis of the cytoplasmic compartment, leaving a single intact nucleus in each microwell^16–18^. Organelles can also be captured using antibodies against proteins common on the surface^19^ or using centrifugal based microfluidic devices that can separate cell debris from mitochondria^20,21^. The remaining intact nucleus or organelles are subsequently subjected to genomic analyses, chromatin studies, or nuclear protein assays.

Here, we sought to understand if we could develop a single-cell western blot (scWB) optimized to resolve the cytoplasmic CK8 proteoforms while keeping each cell’s nucleus intact. For this, after evaluating the presence of CK8 proteoforms in MCF7 cells via mass spectrometry, we evaluated several detergents for electromigration and nuclear stability in three breast cancer cell lines with different CK8 expression (MCF7, SKBR3, and MDA-MB-231). We then evaluated electrophoresis conditions to achieve a separation resolution that enables proteoform identification.

## Results and discussion

### Separation of CK8 proteoforms is possible using whole-cell lysis buffers

We first evaluated the utility of scWB to resolve CK8 proteoforms, with and without an intact nucleus in MCF7 cells (**Fig. 1A**). We performed scWB with a whole-cell lysis buffer (RIPA)^22^. The RIPA cell-lysis buffer is supplemented with a 2% w/v sodium dodecyl sulfate (SDS) content for solubilizing both cytoplasmic and nuclear membranes and linearizing cellular proteins for mass-based electrophoresis. Using RIPA, we could solubilize CK8, which exhibited detectable proteoforms under these conditions. We observe that 10.3% of the single cells present a second proteoform (n=629, **Fig. 1B, D**).

**Figure 1.**
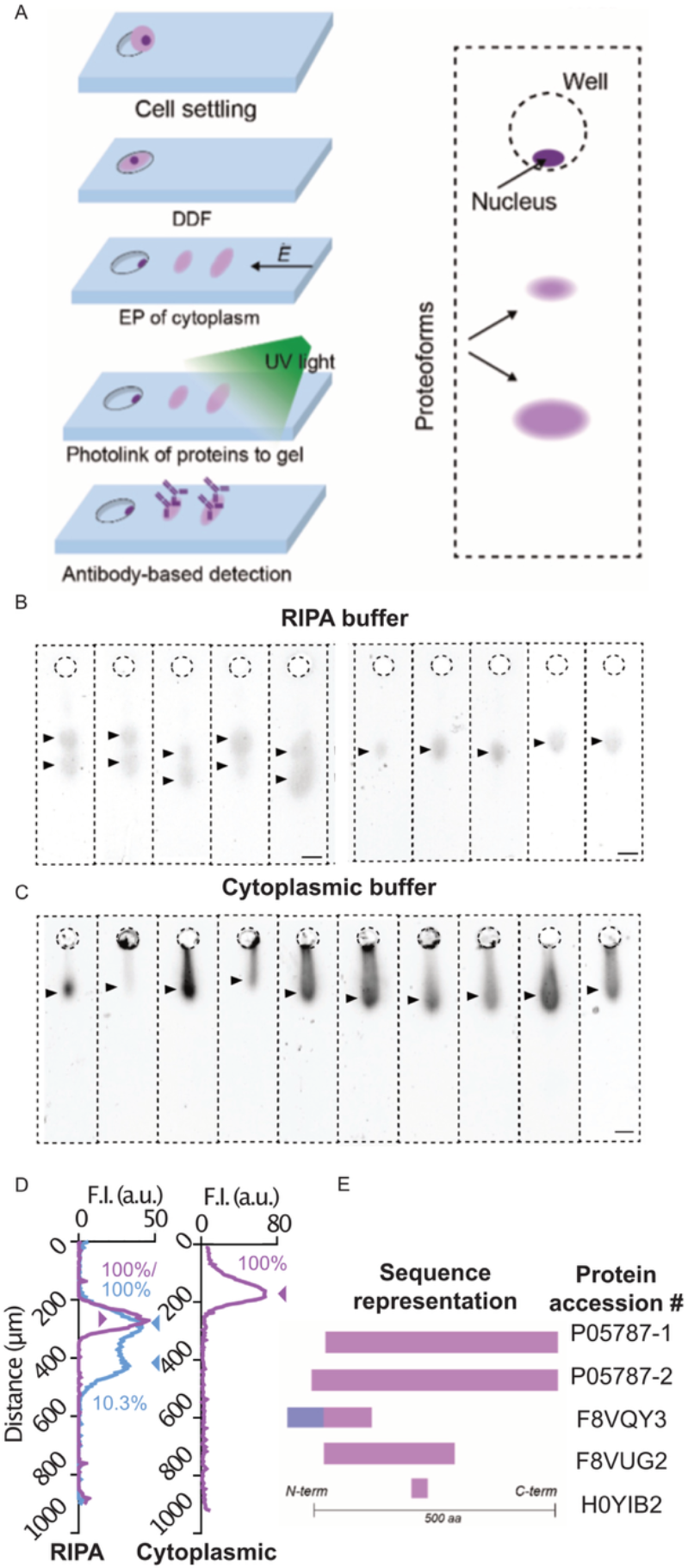
CK8 proteoforms presence in MCF7 cells. A) (Left panel) Workflow schematic for scWB with differential detergent fractionation (DDF). Steps include: gravity-based cell settling into microwells, DDF lysis settled cells electrophoresis (EP) to separate proteoforms by molecular mass with subsequent light-based immobilization of protein to the benzophenone-methacrylate containing polyacrylamide gel, and antibody-based probing for fluorescence detection. (Right panel) Schematic of the desired result, with the nucleus contained in the microwell and the proteoforms separated in the polyacrylamide gel. B) Examples of scWB protein bands after dissolution with RIPA buffer, which solubilizes both nuclear and cytoplasmic proteins, showing the presence of the main protein and a proteoform (left panel) or only the main protein (right panel). C) Examples of scWB after partial solubilization achieved using the previously reported cytoplasmic buffer^16^, which solubilizes only the cytoplasmic compartment. The trail shows partially solubilized filaments of CK8. D) Graph representing the grey scale intensity profile of a scWB band for RIPA and cytoplasmic buffers. The RIPA-lysed profile presents two curves, a purple that represents a single form, and a blue that shows the presence of the main form and a proteoform. In the cytoplasmic-lysed profile, only a single form (purple) is detected. 100% of the cells presented the main form, but only 10.3% presented the proteoform in the RIPA-lysed cells (n=629 for RIPA, n=800 for cytoplasmic). Scale bar: 30 μm. E) Schematic representing the proteoforms detected by top-down mass spectrometry in a bulk MCF7 cell suspension. Regions of sequence alignment identified by the Clustal Omega program from UniProt are highlighted in light purple, while dark purple indicates distinct sequences.

To retain the nuclear compartment while assessing CK8 proteoforms, we performed DDF scWB and assessed a cytoplasmic buffer previously developed by our team for single-cell DDF for scWB^16–18^,consisting of Triton X-100, digitonin, and Tris-glycine. With these conditions, we did not observe the CK8 proteoform in a population of 800 cells (**Fig. 1C, D**). There are two primary explanations for this: a) the proteoform does not properly solubilize with the cytoplasmic buffer, likely remaining interlinked together with the main proteoform, or b) the migration of both proteoforms is overlapping, as this native buffer does not contain SDS, thus preserving native protein structure and protein charge state. This finding suggests that advancing single-cell CK proteoform analysis while maintaining nuclear integrity will require the development of new chemical solubilization strategies, thus forming the rationale for the present study.

To confirm the presence of CK8 proteoforms in MCF7 and validate our results, we performed top-down mass spectrometry. We identified CK8 proteoforms accession #P057878-1, #P05787-2, #F8VQY3, #F8VUG2, and #H0YUB2 (**Fig. 1E** and **SI Table 2**). These findings indicate that the MCF7 cells express both confirmed alternative-splicing CK8 isoforms, as well as additional CK8 fragments.

Based on the electrophoretic migration distance observed in the scWB, the main CK8 proteoform observed is posited to be the main CK8 form or isoform (53.7 and 56.6 kDa, respectively). However, the resolving capability of scWB is not expected to resolve such small differences. This difference is 2.9 kDa, which constitutes < 12% of molecular mass, reported as the smallest resolvable mass difference to date for scWB^23^.Thus, we attribute the other detected proteoform to the fragment #F8VUG2 (30.9 kDa), observed in MCF7 cells via mass spectrometry.

### Nonionic detergents enhance solubilization and maintain nuclear stability

The composition of the detergent cocktail in any DDF lysis buffer impacts protein solubilization, especially at the single-cell level. The scWB demands protein solubilization in just a few seconds, because the open microwell format means that diffusion of the concomitantly solubilizing proteins out of the microwell curtails detection sensitivity of the completed scWB^24^. Thus, solubilization times of 100-1000’s of seconds are not possible or require alternative strategies, such as droplet encapsulation^25^, making the choice of detergent critical to achieve proper protein solubilization. Detergents result in different solubilization efficiencies for each protein species, a product of each protein distinct properties, including charges, hydrophilic and hydrophobic regions, and 3D conformation. Buffers are typically composed of a combination of three commonly used detergent types: anionic, zwitterionic, and nonionic detergents. Anionic detergents, such as SDS, generally destabilize all cellular membranes and thus are not compatible with cell fractionation (i.e., the whole cell lyses)^26^. Other types of detergents, including zwitterionic and nonionic are typically mild and can break lipid-lipid and lipid-protein interactions, but preserve protein-protein interactions intact, or even are biologically active^23^.

Here, we evaluated several detergents to determine the efficacy in solubilizing CK8 proteoforms (**SI Fig. 1**). We used the cytoplasmic buffer as the base composition and tested the addition of Brij-35, n-dodecacyl-β-maltosidase (DDM), IGEPAL CA- 630, and urea. **Fig. 2A** shows the migration distances of CK8 achieved with each of the buffers. While lysis with urea, a chaotropic agent, presents one of the highest migrations, it was not further used due to the damage it produces to the nucleus (**Fig. 2B, right panel**), causing DNA to migrate into the gel. This was not observed in buffers based on nonionic detergents. Given their inability to dissolve protein-protein interactions, nonionic detergents cannot dissolve the nuclear lamina structure and consequently maintain nuclei integrity in a manner that makes said nuclei suitable for further analysis (**Fig. 2B, left panel**).

**Figure 2.**
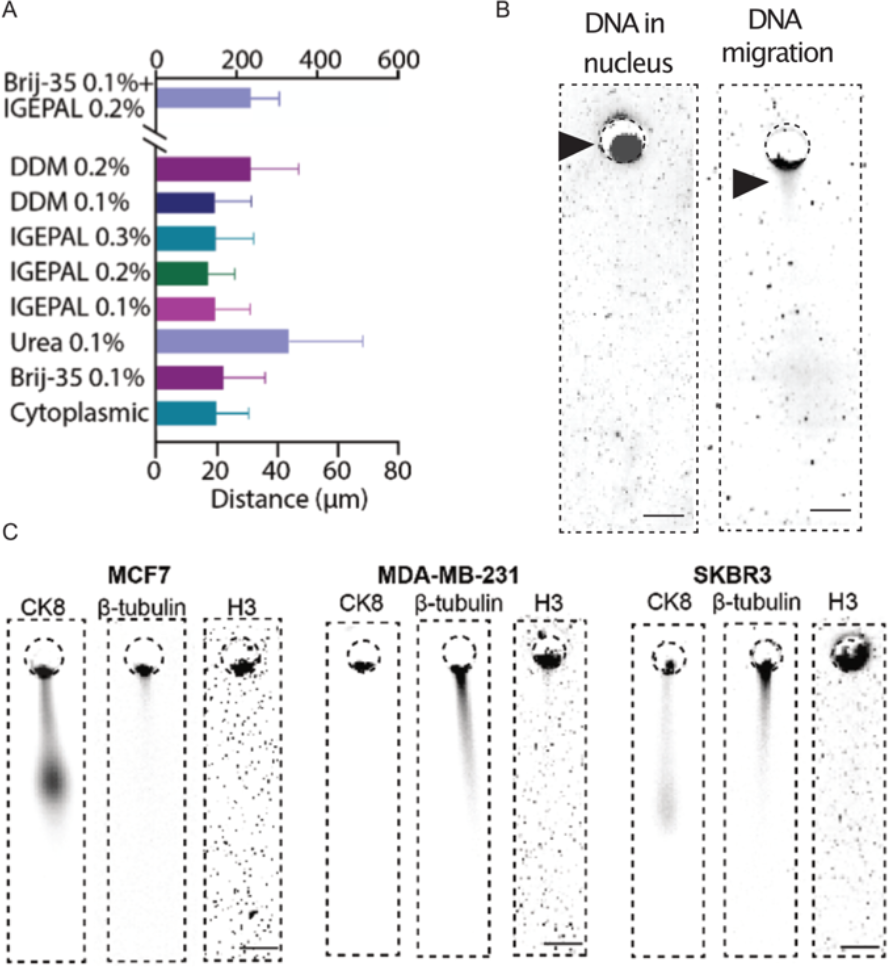
Effects of the solubilization buffer on CK8 migration and nuclear integrity. A) Bar plot showing CK8 migration distance into the polyacrylamide gel. The longest migration distance was observed with Brij-35 0.1%+IGEPAL 0.2% buffer (DDF buffer) and 0.1% urea buffer. B) Micrograph representing a microwell stained for DNA using SYBR gold after scWB. The left image shows a preserved nucleus (use of DDF buffer), while the right image has fragmented DNA that penetrates into the gel (use of urea 0.1% buffer). C) Micrographs reporting DDF lysis of representative single cells from three breast cancer cell lines MCF7, MDA-MB-231, and SKBR3, cytoplasmic proteins CK8 and β-tubulin, which migrate into the gel, while histone H3, a nuclear marker intercalated in the DNA, does not inject into the gel. Image gain was substantially increased for imaging in H3 vs other markers to ensure that no migration of protein is observed, and thus noise is observed in these micrographs. Scale bar: 30 μm.

Neither of the nonionic detergents alone generated a sufficient differential electromigration to resolve CK8 forms. However, a combination of Brij-35 0.1% + IGEPAL 0.2% (DDF buffer) had a much higher migration distance than any of the nonionic detergents alone (**Fig. 2A**). After solubilization with either of the buffers, we could still observe remnant CK8 inside the nucleus. This could be explained by the presence of CK8 in the nucleus, but also by partial solubilization of protein.

CK8 detection by electrophoresis is observed in 34% of microwells containing cells (**SI Fig. 1B**), in contrast to β-tubulin detection, which reached up to a 68% solubilization under some of the conditions (**SI Fig. 1D**). Despite the higher solubilization and the stronger signal observed in β-tubulin, the solubilization of β-tubulin protein with either of the buffers was not complete. Protein species require individualized approaches for solubilization^27^, thus precluding development of a “universal buffer” that can solubilize most of a diverse physicochemical universe of protein molecules. The electrophoretic migration distance from the protein injection point (i.e., the microwell lip) was measured to be non-uniform across the different buffers (**Fig. 1A**), suggesting that different nonionic detergents facilitate proteoform electrophoretic separation by exploiting distinct migration patterns related to molecular mass, surface charge, and structure.

### Nuclear stability after DDF and electrophoresis depends on cell line

For a DDF protocol to be effective, the physical and chemical integrity of each nucleus needs to be preserved. While nonionic detergents are mild and preserve most protein-protein interactions, this effect could vary across different cell lines. **Fig. 2C** illustrates the migration of CK8 and β-tubulin in three different breast cancer cell lines (MCF7, SKBR3, and MDA-MB-231), confirming the solubilization of proteins under the selected conditions. The migration of CK8 corresponds to the expected values, with no CK8 detected in the basal cell line, MDA-MB-231, while MCF7, a luminal cell line, and SKBR3, a HER2 overexpressing cell line, showed migration^16^. In particular, we observed the separated CK8 proteoforms in MCF7 but not SKBR3.

To assess nuclear stability, we performed staining with histone H3 as a marker. Histones are an integral part of the chromatin structure, playing a key role in DNA condensation. The absence of histone migration into the gel is used as a proxy for an intact nucleus. Experimentally, we observe that H3 remains retained within the microwell, suggesting preserved nuclear stability^28^.

For comparison, **SI Fig. 2** depicts a disrupted nucleus from an MCF7 cell, where the nuclear content is migrating inside of the gel, and thus the nuclear structure is compromised. This highlights the importance of matching cell lysis buffer composition to cell type in achieving reliable nuclear preservation in DDF.

### Proteoforms of CK8 can be resolved with the DDF buffer

From the conditions considered, we resolved CK8 proteoforms in two buffers: Brij-35 0.1%, and IGEPAL 0.2%, which led us to combine both detergents to increase proteoform solubilization. Using a combined buffer composition (Brij-35 0.1%+IGEPAL 0.2%) increased the percentage of cells with detectable proteoforms to 14.3% (**Fig. 3A**), considerably more than using either detergent alone and comparable to results with the RIPA buffer. However, Brij-35 has the lowest CK8 solubilization efficiency across the nonionic detergents tested, and its presence in the DDF buffer reduces the total CK8 solubilization when compared to the IGEPAL 0.2%. This is not the case for β-tubulin, which exhibits poor solubilization with DDM, but the highest with the DDF buffer (**SI Fig. 1**).

**Figure 3.**
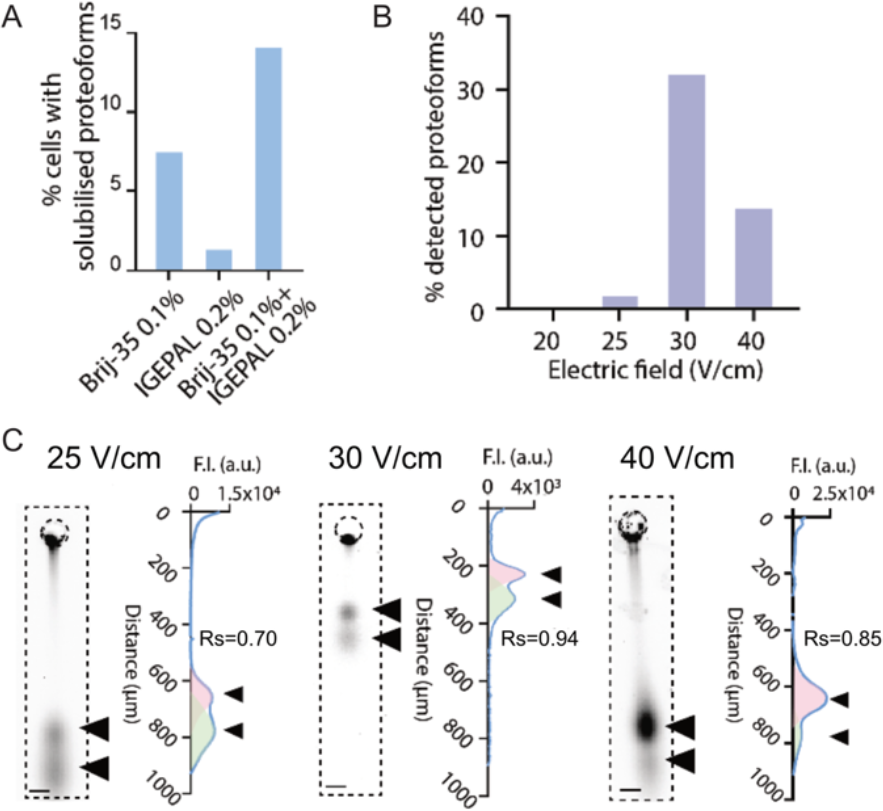
CK8 proteoform separation on scWBs. A) Percentage of cells with detectable proteoforms per buffer where proteoforms were visualized (total cells with migration into the gel n=26 for Brij-35 0.1%, n=132 for IGEPAL 0.2%, and n=35 for Brij-35 0.1%+IGEPAL 0.2%). B) Percentage of cells with detectable proteoforms across a set of applied electric field conditions. C) Plot showing the separation resolution of CK8 proteoforms across a set of applied electric field conditions, with the corresponding micrograph showing the main form and the proteoform. Electrophoresis times were adjusted to 70, 50, and 40 s for 25 to 40 V/cm, respectively, as the same electrophoresis time would not separate the proteoforms under all conditions. The triangles indicate the location of each proteoform in the electropherogram.

In order to sufficiently resolve the detectable CK8 proteoforms, various parameters such as lysis time, buffer temperature, and electric field were evaluated. In all cases, adjusting the parameters involved trade-offs. For CK8, lower solubilization temperatures (55°C) offered significantly higher signal intensity per area (**SI Fig. 3**); however, solubilization and separation resolution were improved at 85°C. Furthermore, the migration distance of CK8 increased with temperature, likely due to a better solubilization of CK8 filaments (**SI Fig. 3A, C**). The lysis time did not show a clear correlation with the number of solubilized cells, although 30 s solubilization gave generally the best intensity/area ratio for CK8 (**SI Fig. 3B**). In the case of β-tubulin, 20 s met our performance goals based on the median intensity/area of the cells, while 75°C was the optimal lysis temperature (**SI Fig. 3C-D**).

To maximize the CK8 proteoform separation resolution, we analyzed the electrophoretic mobility of the two CK8 proteoforms resolved by scWB using Ferguson plots with varying polyacrylamide-gel concentrations (%T) (**SI Fig. 4-5**). Both CK8 proteoforms show similar values for the y-intercept (yfull-length=-15x-2.66 and yfragment=-13.0x-2.64), suggesting that the use of different total acrylamide (%T) concentrations in the separation gel will be unlikely to enhance separation resolution. In addition, the non- linearity of the relationship suggests a non-spherical conformation for the CK8 molecules, consistent with reports of CK8 being a filamentous protein^29^.

Sweeping a range of applied electric field conditions, however, suggested that this parameter offered the largest impact on the separation resolution of the proteoforms (**Fig. 3C**). At 30 V/cm, separation resolution was the largest (Rs=0.94, calculated as *Rs=2(d2-d1)/(w1+w2)*, where *d* is the distance to the peak, and *w* is the peak width) and the highest number of proteoforms detected as a percentage of total electrophoresed cells. An applied electric field of 25 V/cm gave an Rs=0.7, whereas 40 V/cm yielded Rs=0.85. Conversely, at 20 V/cm proteoforms were not resolvable, and overall band intensity was weak. The electrical field also impacted the number of detected proteoforms, with 30 V/cm showing the highest percentage (**Fig. 3B**). In cases of poor separation resolution, the presence of CK8 proteoforms can only be identified by a skewed Gaussian peak in comparison with a compact circular band observed in cells with only one proteoform.

## Experimental

### SU-8 mold fabrication

Molds for soft lithography were fabricated as previously described^22^. A layer of SU-8 3050 was spun over a silicon wafer for 10 s at 500 rpm with an acceleration of 100 rpm, and then for 30 s at 4000 rpm with an acceleration of 300 rpm to achieve a layer thickness of 40 μm. The wafer was soft baked for 2 min at 65°C, 15 min at 95°C and then 3 min at 65°C. The wafer was exposed to a UV dose of 385 mJ/cm^2^ and then baked post-exposure for 1 min 65°C, 5 min at 95°C and then 1 min at 65°C. The wafer was developed for 5 min on a shaker and then hard baked at 200°C for 20 min.

### Single-cell western blot

The scWB was performed as previously reported^22^. Briefly, gels were fabricated mixing 7% acrylamide:bis-acrylamide (29:1, Sigma-Aldrich), 3 mM BPMA (BP-APMA, custom synthesis by Raybow Pharmaceutical), 0.08% ammonium persulfate (Sigma Aldrich), and 0.08% tetramethylethylenediamine (Sigma Aldrich). The solution was then incorporated between a wafer and a silanized glass slide^22^ until the end of polymerization after 15 min, when the gel was detached from the wafer and deposited in PBS for 1h. The gel was then dried under a nitrogen stream, and 250 μL pf 1 million/mL of strained cells were deposited on the gel for cell settling. After 10 min, the excess cells were removed with PBS, and the gels were placed in the Milo system (Protein Simple, Bio-Techne) for electrophoresis. The lysis and electrophoresis times, and the electric field were adjusted on an experimental basis, with lysis 30 s and electrophoresis 30 s unless otherwise noted. The photoactivation time was set to 45 s. After the electrophoresis step, the gel was deposited in 1x Tris Buffered Saline + 0.1% Tween-20 (TBST) for 1h. The gel was then incubated with primary antibody for 1 h, washed in TBST for 45 min with one wash exchange after 20 min, and incubated with the secondary antibody for 1h. The sample was washed in TBST for 45 min with one wash exchange and then rinsed in water to remove salts. The gel was dried under a nitrogen stream and imaged using Genepix Microarray Scanner (Genepix 4300A, Molecular Devices), with a resolution of 5 μm. The channel 535 used a laser power of 90% and a gain of 500, and channel 635 used a laser power of 70% and a gain of 700.

Primary antibodies were β-tubulin (100 μg/mL, ab6046 Abcam), CK8 (2.5 μg/mL, ab9023 Abcam), histone H3 (100 μg/mL, ab1791 Abcam) and secondary antibodies were anti-rabbit-633 (A21071 Thermo Fisher) and anti-mouse-532 (A11002 Thermo Fisher), all at 100 μg/mL. DNA was stained using SYBR gold at the suggested concentration (S11494 Thermo Fisher).

Data analysis was performed using a Python custom algorithm based on ref.^30^, using numpy, pandas, tkinter, and pillow. Briefly, the images were cut into single bands, a thresholding function was used for segmentation of the band, and the intensity was extracted.

Plotting was performed using Prism. Statistics were performed using the Kruskal-Wallis test for non-parametric datasets. The scWB micrographs presented here are inverted grayscale and contrast-adjusted for visualization.

### Buffer composition

The RIPA buffer composition used was 1% w/v SDS, 0.5% w/v sodium deoxycholate, 0.1% v/v Triton-X100, and 0.5x of Tris Glycine. The cytoplasmic base buffer composition was 1% v/v Triton X-100, 0.125 mg/mL digitonin 99% pure, and 0.5x Tris Glycine. Buffers from Fig. 2 were formulated by adding the described w/v (DDM, urea (both Sigma Aldrich)) or v/v (Brij-35 (Thermo Fisher), IGEPAL (Abcam)) of the detergent in the cytoplasmic base buffer (**SI Table 1**).

### Top-down mass spectrometry

The fractions of proteins were obtained using the protocol described in PEPPI-MS^31^. The fractions (6 μL) were analyzed by online capillary nanoLC-MS/MS using a 40 cm reversed phase column fabricated in-house (50 µm inner diameter, packed with ReproSil-Gold C8-3 μm resin (Dr. Maisch GmbH)) that was equipped with a laser-pulled nanoelectrospray emitter tip. Peptides were eluted at a flow rate of 100 nL/min using a linear gradient of 2–40% buffer B in 140 min (buffer A: 0.05% Formic acid in water; buffer B: 0.05% Formic acid and 95% acetonitrile in water) in an Thermo Fisher Easy-nLC1200 nanoLC system. Peptides were ionized using a FLEX ion source (Thermo Fisher) using electrospray ionization into an Fusion Lumos Tribrid Orbitrap Mass Spectrometer (Thermo Fisher Scientific). Data was acquired in orbi-trap mode. Mass spectrometry data analysis was performed using TopPIC^32^ and TopMSV^33^. The E-value of the detected proteoforms is ∼10^−3^-10^−4^. The summary of the results is provided in **SI Table 2**. The reported proteoforms were identified in two biological replicates of PEPPI-MS. Proteoforms identified in only one of the replicates were discarded.

## Conclusions

The diversity of physicochemical properties of protein molecules demands customization and optimization of lysis conditions to achieve an effective protein separation^34,35^, especially at the single-cell level. Such optimization is not uncommon and parallels development of crystallography assays, where obtaining protein crystals requires extensive testing of different conditions. Similarly, successful selective lysis and proteoform separation in electrophoretic gels will depend on the specific conditions employed. Such selectivity is not required for scWB performed with whole cell solubilization, i.e., no nuclear isolation.

Here, we report assay development for the solubilization and detection of CK8 proteoforms while preserving nuclear integrity in single breast cancer cells. The proper solubilization of filamentous proteins while keeping the nuclear lamina intact is a particularly difficult assay development challenge, because the same conditions that solubilize the protein-protein interactions also may solubilize the nuclear lamina, thus obstructing DDF. By selecting nonionic detergents and adjusting electrophoretic conditions, we obtain enhanced separation and, thus, detection of specific CK8 proteoforms across different cell lines. We achieve a separation resolution of 0.94, higher than previously reported^18^ and, provide an assay for CK8 proteoforms where other immunoassay-based single-cell tools like flow cytometry fall short.

Owning to the single-cell resolution of the western blot, our approach provides a platform to investigate the biological heterogeneity of CK8 fragments while retaining the nuclear compartment separate. While ultimate confirmation of proteoform identity benefits from mass spectrometry, we observe that both whole-cell lysis (RIPA) and the cytoplasmic lysis (DDF) yields two CK8 peaks. The main CK8 forms have comparable molecular mass and a pI corresponding to the alternative spliced forms (molecular mass=53.7 and 56.6 kDa, and pI=5.52 and 5.37 as calculated by the ExPASy Compute pI/Mw tool^36^). Thus, the observed proteoform most likely corresponds to #F8VUG2 (30.9 kDa). Although these alternative splicing products and protein fragments have been linked to distinct disease states, their precise biological significance remains largely unexplored.

The separation of cytoplasmic compartments enabled by DDF is in turn the first step for a post-hoc analysis of nuclear proteins and different omics, such as transcriptome, genome, or epigenome, requiring the integration of DDF with other techniques^17,18^. CK8 presents several transcript products of alternative splicing, but the protein also shows protein variability product of protease activity.

While the DDF scWB presented here enables the examination of many of these possibilities, it still presents some limitations. For instance, the separation of closely sized spliced proteoforms remains a challenge, and the percentage of solubilized proteins remains low in all tested conditions. Nonetheless, this work represents a step forward in aiding single-cell proteoform analysis, enabling protein characterization while preserving the integrity of cellular compartments for downstream multi-omic investigations.

## Supporting information

Supplementary Table 2. Mass spectrometry protein analysis

## Data availability

The data supporting this article have been included as part of the Supplementary Information. Mass spectrometry data for this article are available at ProteomeXchange with identifier PXD063523.

## Acknowledgements

A.E.H. acknowledges financial support from the Investigator Program of the Chan Zuckerberg Biohub San Francisco and the US National Institutes of Health (NIH R01CA20301). A.F.K. acknowledges support from the Swiss National Science Foundation (SNSF) Postdoc Mobility fellowship (P500PN_214236). The photolithography was performed in the QB3 Biomolecular Nanotechnology Center and mass spectrometry was performed by the QB3 Vincent J. Coates Proteomics/Mass Spectrometry Laboratory by Dr. Robert Maxwell, both at UC Berkeley.

