## Supplementary information for "Single-cell western blotting of cytoplasmic cytokeratin 8 proteoforms"

**SI Figure 1. The protein CK8 and β-tubulin (control) for nonionic buffer testing development**.

**SI Figure 2. Stability of the nucleus is cell-line dependent.**

**SI Figure 3. Lysis buffer temperature and lysis time impact on CK8 and β-tubulin solubilization**.

**SI Figure 4. Ferguson analysis of CK8 proteoforms.**

**SI Figure 5. Influence of the percentage of acrylamide in proteoform migration.**

**SI Table 1. Composition of employed buffers.**

**SI Figure 1. The protein CK8 and β-tubulin (control) for nonionic buffer testing development**. A) Violin plot representing intensity profiles normalized by the total area of the protein spot for CK8 across tested buffers. The number of microwells with cells in the scWB ranges from 14 to 124. B) Bar plot representing the percentage of solubilized cells using different detergents. (A) Signal intensity and (B) solubilization percentage of β-tubulin under different detergent conditions on MCF7 cells. Lysis time was 30s and electrophoresis time was 30s, gels were 7%T and the buffer was at 55°C. The number of microwells with cells in the scWB ranges from 212 to 454.

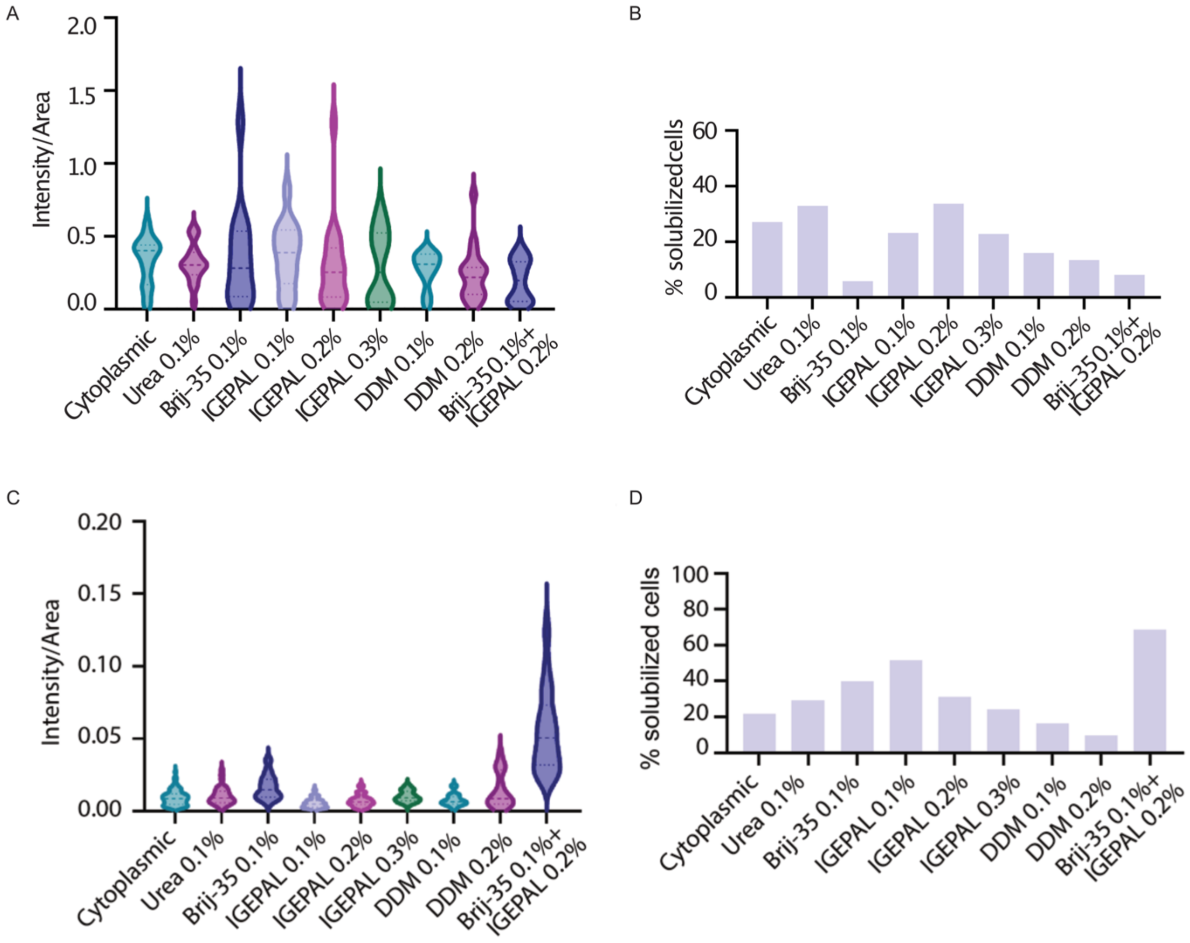

**SI Figure 2. Stability of the nucleus is cell-line dependent.** Transfected MCF7 cells showed migration of histone H3 into the gel after applying the same protocol as regular MCF7s, which highlights the need to perform buffer optimization for each new cell line. For comparison, the right panel shows a full solubilization of H3 using the RIPA buffer. The black triangles indicate the location of the fluorescent signal. Scale bar: 30 µm.

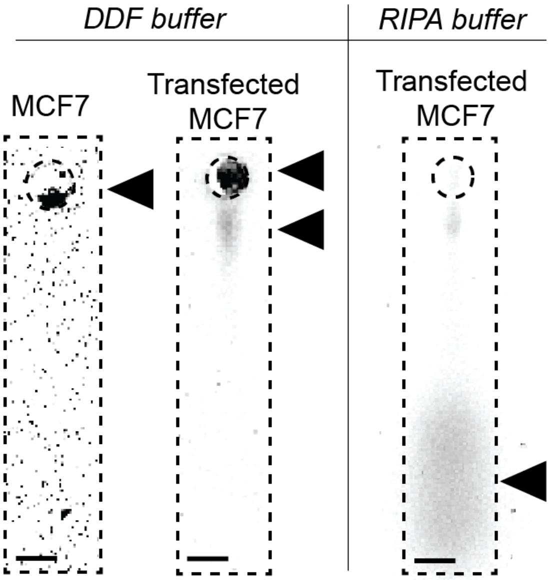

**SI Figure 3. Lysis buffer temperature and lysis time impact on CK8 and β-tubulin solubilization.** (A) Optimization of buffer temperature for CK8. Lysis time was 30s and electrophoresis time was 30 s, gels were 7%T. For each temperature, n=78, 31, 78, and 40, respectively. (B) Plot showing the migration distance of the CK8 bands with increasing temperatures. The right panel shows that the solubilization of the protein is better at higher temperatures. Scale bar: 30 μm. (C) Optimization of buffer temperature for β-tubulin. Lysis time was 30s and electrophoresis time was 30s, gels were 7%T. For β-tubulin, n=349, 363, and 286, respectively. (D) Optimization of buffer lysis time for β-tubulin. Electrophoresis time was 30s, gels were 7%T, and the buffer temperature was 55°C. For β-tubulin, n=214, 349, 468, 120, and 113, respectively. Significance is as follows: * P ≤ 0.05, ** P ≤ 0.01, *** P ≤ 0.001, **** P ≤ 0.0001. (E) Optimization of buffer lysis time for CK8. Electrophoresis time was 30s, gels were 7%T, and the buffer temperature was 55°C. For each lysis time, n=134, 78, 153, 56, and 29, respectively. Significance is as follows: * P ≤ 0.05, ** P≤ 0.01, *** P ≤ 0.001, **** P ≤ 0.0001.

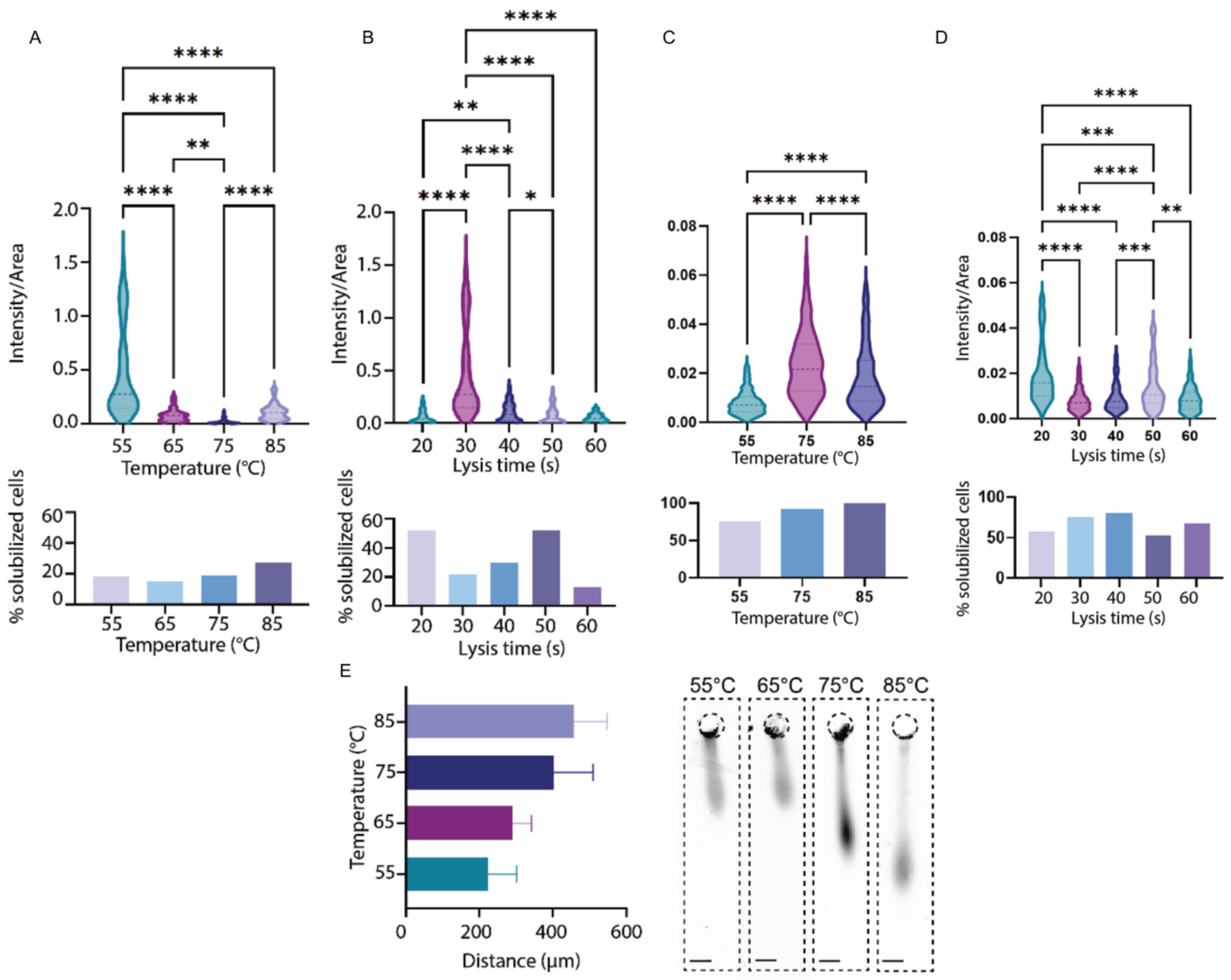

**SI Figure 4. Ferguson analysis of CK8 proteoforms**. Main CK8 form is at left, and proteoform is at right. Applied E = 30 V/cm.

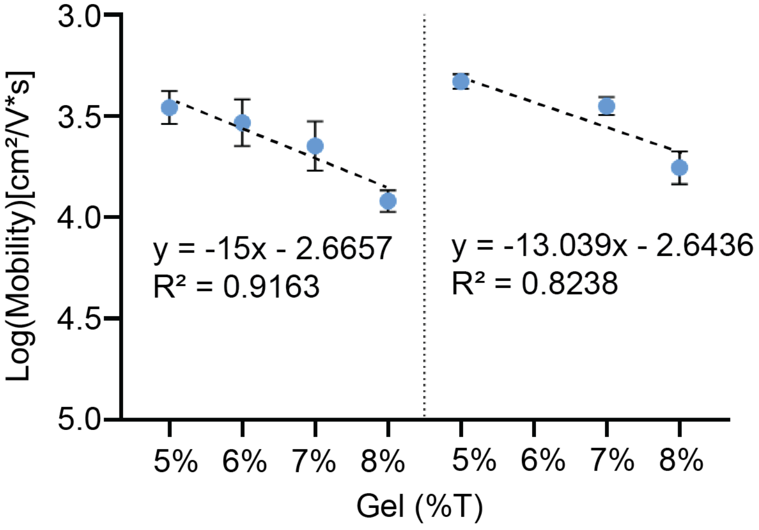

**SI Figure 5. Influence of the percentage of acrylamide in proteoform migration.** Micrograph matrix showing proteoforms across two electric fields and four %T.

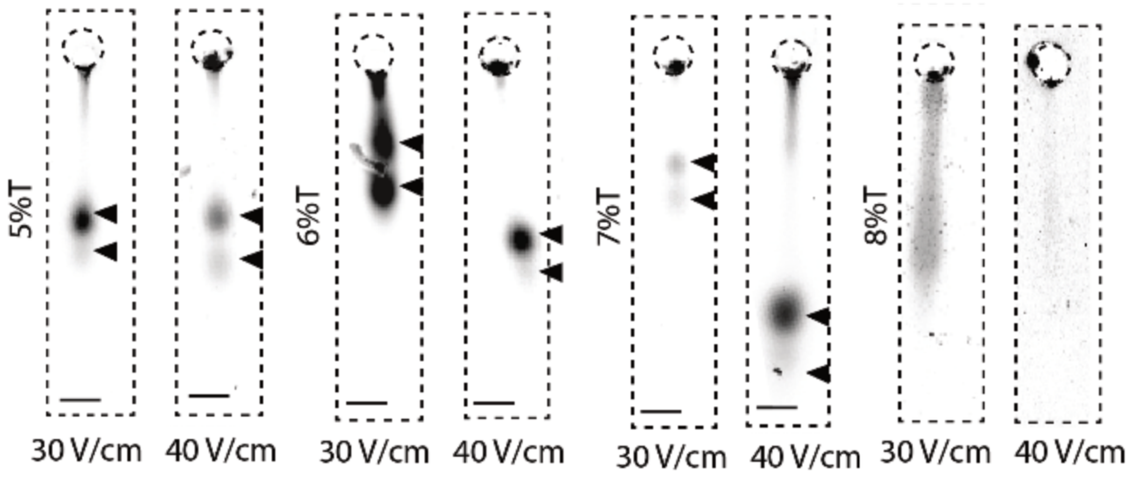

### SI Table 1. Composition of employed buffers.

#### Cytoplasmic buffer

| Chemical | Final concentration | Amount |
|---|---|---|
| Triton X-100 | 1% v/v | 300 µL |
| Tris-Glycine | 0.5x | 1.5 mL |
| Digitonin | 0.125 mg/mL | 3.75 mg |
| Water | | 28.2 mL |

#### DDF buffers

| Chemical | Final concentration | Amount |
|---|---|---|
| Triton X-100 | 1% v/v | 300 µL |
| Tris-Glycine | 0.5x | 1.5 mL |
| Digitonin | 0.125 mg/mL | 3.75 mg |
| IGEPAL | 0.1% v/v | 300 µL |
| Water | | 27.9 mL |

| Chemical | Final concentration | Amount |
|---|---|---|
| Triton X-100 | 1% v/v | 300 µL |
| Tris-Glycine | 0.5x | 1.5 mL |
| Digitonin | 0.125 mg/mL | 3.75 mg |
| Brij-35 | 0.1% v/v | 300 µL |
| Water | | 27.9 mL |

| Chemical | Final concentration | Amount |
|---|---|---|
| Triton X-100 | 1% v/v | 300 µL |
| Tris-Glycine | 0.5x | 1.5 mL |
| Digitonin | 0.125 mg/mL | 3.75 mg |
| IGEPAL | 0.2% v/v | 600 µL |
| Water | | 27.6 mL |

| Chemical | Final concentration | Amount |
|---|---|---|
| Triton X-100 | 1% v/v | 300 µL |
| Tris-Glycine | 0.5x | 1.5 mL |
| Digitonin | 0.125 mg/mL | 3.75 mg |
| Urea | 0.1% w/v | 30 mg |
| Water | | 28.2 mL |

| Chemical | Final concentration | Amount |
|---|---|---|
| Triton X-100 | 1% v/v | 300 µL |
| Tris-Glycine | 0.5x | 1.5 mL |
| Digitonin | 0.125 mg/mL | 3.75 mg |
| IGEPAL | 0.3% v/v | 900 µL |
| Water | | 27.3 mL |

| Chemical | Final concentration | Amount |
|---|---|---|
| Triton X-100 | 1% v/v | 300 µL |
| Tris-Glycine | 0.5x | 1.5 mL |
| Digitonin | 0.125 mg/mL | 3.75 mg |
| Brij-35 | 0.1% v/v | 300 µL |
| IGEPAL | 0.2% v/v | 600 µL |
| Water | | 27.3 mL |

| Chemical | Final concentration | Amount |
|---|---|---|
| Triton X-100 | 1% v/v | 300 µL |
| Tris-Glycine | 0.5x | 1.5 mL |
| Digitonin | 0.125 mg/mL | 3.75 mg |
| DDM | 0.1% w/v | 30 mg |
| Water | | 28.2 mL |

| Chemical | Final concentration | Amount |
|---|---|---|
| Triton X-100 | 1% v/v | 300 µL |
| Tris-Glycine | 0.5x | 1.5 mL |
| Digitonin | 0.125 mg/mL | 3.75 mg |
| DDM | 0.2% w/v | 60 mg |
| Water | | 28.2 mL |
